# Maternal proteins that are phospho-regulated upon egg activation include crucial factors for oogenesis, egg activation and embryogenesis in *Drosophila melanogaster*

**DOI:** 10.1101/218925

**Authors:** Zijing Zhang, Amber R. Krauchunas, Stephanie Huang, Mariana F. Wolfner

## Abstract

Egg activation is essential for the successful transition from a mature oocyte to a developmentally competent egg. It consists of a series of events including the resumption and completion of meiosis, initiation of translation of some maternal mRNAs and destruction of others, and changes to the vitelline envelope. This drastic change of cell state is accompanied by large scale alteration of the phospho-proteome of the cell. Despite the importance of this transition in cell and developmental state, it has been difficult to find many of its regulators. We hypothesize that phosphorylation state changes between oocyte and early embryo regulate the activities of proteins that are necessary during or after this transition, and thus that the set of phospho-regulated proteins would be an enriched source for finding critical players in the egg-to-embryo transition. To test this, we used germline-specific RNAi to examine the function of 189 maternal proteins that are phospho-regulated during egg activation in *Drosophila melanogaster*. We identified 53 genes whose knockdown reduced or abolished egg production, as well as 50 genes for which maternal knockdown led to significant impairment or abolishment of the eggs’ ability to hatch (hatchability). We observed different stages of developmental arrest in the embryos with impaired hatchability and several distinct categories of abnormalities in the maternal knockdown embryos that arrest early in development, suggesting potential roles of the candidates in egg activation and early embryogenesis. Our results, validated by our detection of multiple genes with previously-documented maternal-effect phenotypes among the proteins we tested, revealed 15 genes with new roles in egg activation and early embryogenesis. Given that protein phospho-regulation also occurs during this transition in other organisms, we suggest that the phospho-regulated proteins may provide an enriched dataset for identifying important players in the egg-to-embryo transition.

## INTRODUCTION

At the end of oogenesis, mature oocytes arrest in meiosis with repressed transcription and a rich maternal deposit of mRNAs and proteins. For development to proceed, the oocyte needs to restart quiescent cellular activities. This occurs through the process of egg activation, which consists of a series of cellular events including the resumption and completion of meiosis, modifications of the outer egg coverings, and dynamic regulations of maternal mRNAs (Horner & Wolfner 2008b, Krauchunas & Wolfner 2013, Laver et al 2015, Machaty et al 2017, Marcello & Singson 2010, Von Stetina & Orr-Weaver 2011). Egg activation is an intricate process through which the oocyte shifts gears from oogenesis to embryogenesis. It marks the transition during which a differentiated gamete transforms into a totipotent zygote that gives rise to all the cell types of the adult. The regulatory mechanisms of this transition are vital for female fertility. Although some of these factors have been identified genetically (Cui et al 2008, Elfring et al 1997, Freeman et al 1986, Lee et al 2003, Lin & Wolfner 1991, Renault et al 2003, Tadros et al 2003), their number is surprisingly small. It seems likely that many molecules essential for this transition were not identified in traditional maternal-effect screens since those would miss genes whose activity is essential for the organism survival. Such molecules might nevertheless play important roles in oogenesis, egg activation, and/or embryogenesis. Here, we show that the proteins that are phospho-modulated during egg activation represent an enriched pool from which to find such regulators.

Egg activation initiates with a rise of Ca^2+^ level in the cytoplasm of the oocyte. This calcium rise is triggered by fertilization in vertebrates and most marine invertebrates (Knott et al 2005, Saunders et al 2002, Steinhardt et al 1977, Yoon et al 2012). In insects, the calcium rise is triggered by the passage of oocyte through the reproductive tract, and it takes place independent of fertilization (Horner & Wolfner 2008a, Kaneuchi et al 2015, Sartain & Wolfner 2013, York-Andersen et al 2015). Since transcription is quiescent during oocyte maturation and egg activation (De Renzis et al 2007, Newport & Kirschner 1982, Zalokar 1976), the dowry of maternal mRNAs and proteins must include all the essential machinery that regulates the cellular events of egg activation in response to the calcium rise. A study in *C. elegans* that compared the oocyte proteome and transcriptome revealed a bias in the composition of the oocyte proteome towards gene products required immediately upon fertilization, whereas the oocyte transcriptome was more highly enriched in genes involved in later embryogenesis (Chik et al 2011). Furthermore, a study in Drosophila found that meiosis can be completed even when translation is inhibited (Page & Orr-Weaver 1997). These findings suggest that the regulation of egg activation, and the early embryonic events that immediately follow, relies heavily on maternally-provided proteins. However, some of these proteins might need to be active at one stage in this transition (e.g. oogenesis) and inactive at another (e.g. early embryogenesis), and thus might be post-translationally modulated to accomplish this.

During egg activation, maternal proteins are dynamically regulated through degradation and post-translational modification (Krauchunas et al 2012, Kronja et al 2014, Laver et al 2015). Among the common types of post-translational modifications, phospho-regulation appears to be especially prevalent at this time. Phosphoproteomic analysis of mature oocyte and fertilized eggs in sea urchin revealed dynamic changes of phosphorylation status at thousands of phosphosites at 2 min and 5 min post fertilization (Guo et al 2015, Roux et al 2006). Similarly, our previous analysis of the phosphoproteome in mature oocytes and activated eggs in *Drosophila melanogaster* uncovered more than 300 maternal proteins whose phosphorylation states changed during egg activation (Krauchunas et al 2012). A functional screen in Drosophila revealed that maternal depletion of 33% of the kinases and 18% of the phosphatases present in early embryo lead to embryogenesis defects (Sopko et al 2014), validating the functional importance of phospho-regulations for egg activation and embryogenesis. In addition, two highly-conserved calcium-dependent regulators essential for egg activation in several species, Ca2+/Calmodulin dependent kinase II (CaMKII) (Xenopus and mouse; (Backs et al 2010, Morin et al 1994)) and calcineurin (Xenopus and Drosophila; (Horner et al 2006, Mochida & Hunt 2007, Takeo et al 2010, Takeo et al 2006)), are a kinase and a phosphatase respectively (Hudmon & Schulman 2002, Rusnak & Mertz 2000). These findings suggest phospho-regulation as a crucial mechanism that modulates activities of maternal proteins needed for the transition from oocyte to early embryo. Consistent with this hypothesis, several maternal proteins that are known to be crucial for egg activation and the onset of syncytial divisions, such as Drosophila Young arrest (YA) and Giant nuclei (GNU), are phospho-regulated during egg activation (Lin & Wolfner 1991, Renault et al 2003, Sackton et al 2009, Yu et al 1999). We hypothesize that maternal proteins that are phospho-regulated during egg activation are enriched in factors crucial for cellular processes before, during, and after egg activation, including ones that function at other time of development.

*Drosophila melanogaster* provides an excellent model for testing this hypothesis. In addition to its utility as a genetic model, Drosophila’s egg activation is independent of fertilization (Horner & Wolfner 2008a), allowing a focus on egg activation without confounding influences of fertilization and subsequent zygotic development. Moreover, genetic tools in Drosophila make it possible to test for gene function specifically in the germline, avoiding problems of lethality in a whole-organism screen. Specifically, the UAS-Gal4 system (Brand & Perrimon 1993) allows robust germline specific RNAi knockdown of target genes (Ni et al 2011) to permit these tests.

Here, we used UAS-Gal4 mediated germline-specific RNAi to screen 189 maternal proteins that are phospho-regulated during egg activation (Krauchunas et al 2012) for their roles in female fertility. We observed maternal-effect phenotypes in the knockdown of 50 genes, 27 of which caused significant arrest in early developmental stages, indicating the vital roles of these genes in the initiation of zygotic development. 15 of these 27 genes have no previously characterized maternal-effects. We also identified 53 genes that are crucial for oogenesis. Our study confirms that the cohort of maternal proteins that are phospho-regulated during egg activation is indeed enriched with factors essential for female fertility, provides evidence that phospho-regulation is a vital regulatory mechanism for egg activation, and identified new genes that are essential for different aspects of female fertility.

## MATERIALS AND METHODS

### Fly stocks

Fly stocks were maintained on standard yeast-glucose-agar media, at 23±2°C, under a 12-hr light-dark cycle.

### Germline specific RNAi

A total of 207 Transgenic RNAi lines (TRiP lines) carrying UAS-shRNA targeting 189 candidates were obtained from the Bloomington stock center, the NIG stock center, and the Vienna Drosophila Resource Center (stock numbers are listed in Table S1) (Perkins et al 2015). 5-10 virgin females from the UAS-shRNA stock were mated to 5-10 males carrying either MTD-Gal4, which drives expression throughout oogenesis (Petrella et al 2007), or matα4-GAL-VP16, which drives expression after the egg chamber exits the germarium stage (Radford et al 2012, Sugimura & Lilly 2006). These crosses generated females that are heterozygous for both UAS-shRNA and the driver construct (referred to here as “RNAi females”). Background-matched control females, heterozygous for the AttP2 (empty insertion site of the UAS-hsRNA construct) and the driver constructs were generated in parallel to serve as controls. RNAi females and control females were both raised and maintained at 27°C.

### Fertility assays

Virgin RNAi females and control females were mated to ORP2 males in single pairs, and mating was confirmed by observation. Males were removed after mating and the females were allowed to lay eggs on standard fly media and were transferred to new culture every 24 hours for 4 days. The number of eggs laid in each vial was counted. The hatch rate was estimated as the proportion of eggs that developed into pupae. For each assay, RNAi females for 5-10 TRiP lines (5-10 RNAi females for each line) and 1 control group (10 control females) were included. Total 4-day egg production and overall embryo hatchability were calculated for each female. The 4-day egg production and embryo hatchability of the 5-10 RNAi females for each TRiP line were compared to that of the 10 control females from the same assay using the student’s T-test (Table S1). RNAi lines that generated fertility phenotypes were re-tested for 2-3 times to ensure reproducibility of the results.

### Reverse-transcription PCR

Stage 14 oocytes were dissected in hypertonic isolation buffer that prevents egg activation (Page & Orr-Weaver 1997). RNA was extracted using TRIzol^®^ (Thermo Fisher). RNA samples were treated with DNase (Promega) and reverse transcribed with SmartScribe Reverse Transcriptase Kits (CloneTech Laboratories, Inc). PCR was performed with cDNA samples and amplification band intensities of knockdown and control samples were compared using ImageJ. The primers used for PCR are listed in Table S2

### Bioinformatics

Peptide sequences were scanned for consensus phosphorylation site using Scansite 2.0 Motif Scan (Obenauer et al 2003). The result included consensus phosphorylation sites identified on all the isoforms of each gene. The enrichment analysis was performed using PANTHER version 10 (Mi et al 2016).

### Immunofluorescence

2-4hr old and 0.5-1.5hr old fertilized embryos were collected on grape juice agar plates, dechorionated with 50% bleach, fixed using heptane/methanol and stained as described in (Horner et al 2006). Mouse anti-tubulin (Sigma, St. Louis, MO, catalog #T5168) was used at a dilution of 1:400. Mouse anti-DROP-1, which stains sperm tails (Karr 1991), (kindly provided by T. Karr at Kyoto University) was used at 1:800 as described in Krauchunas et al 2012. Alexa Fluor^®^ 488 conjugated anti-mouse (Thermo Fisher) was used at 1:200. Propidium Iodide was used at 10 µg/ml. Samples were examined and images were generated using a Leica TCS SP2 confocal microscope at the Cornell Imaging Core.

### Immunoblotting

80-100 oocytes or activated but unfertilized eggs (collected from females mated to the spermless sons of *tudor* females) (Boswell & Mahowald 1985) were homogenized in 10 µl of Protein Extraction Buffer (10 mM Tris, pH 7.5; 20 mM NaF, 2 mM EGTA, 10 mM DTT, 400 nM okadaic acid, and 2% SDS), and the lysate was mixed with 10 µl of SDS loading buffer. Proteins were separated by electrophoresis in 8% polyacrylamide SDS gels. Primary guinea pig anti-Smg antibody (Semotok et al 2005) (kindly provided by H. Lipshitz at University of Toronto) was used at 1:10000. Mouse anti-tubulin was used at dilution of 1:10000 (Sigma, St. Louis, MO, catalog #T5168). Secondary HRP conjugated anti-guinea pig was used at 1:1000 (Jackson Laboratories). Anti-mouse secondary antibody was used at 1:2000 (Jackson Laboratories).

## RESULTS

### Germline specific depletion of phospho-regulated maternal proteins reveals factors crucial for different aspects of female fertility

To test whether the subset of proteins that are phospho-regulated during egg activation are important for female fertility in *Drosophila melanogaster*, we obtained all available TRiP RNAi lines that targeted the phospho-regulated maternal proteins identified in Krauchunas et al 2012 (Krauchunas et al 2012). In the primary screen, a total of 207 TRiP lines, targeting 189 genes, were screened using the MTD-Gal4 driver which drives shRNA production throughout oogenesis (Petrella et al 2007). For the genes whose knockdown led to severely reduced or abolished egg production, we performed a secondary screen using matα4-GAL-VP16, which drives shRNA expression in mid and late oogenesis (Radford et al 2012, Sugimura & Lilly 2006). We expected that for genes that have crucial functions in early oogenesis, the temporal manipulation of RNAi knockdown could allow the depletion of gene products in mature oocytes without severely impacting egg production.

We grouped the 189 genes into six classes according to their knockdown phenotypes in the primary and the secondary screen (Table 1, Figure 1A, Table S1). For the 28 genes in Class 1, knocking down their expression throughout oogenesis significantly reduced or abolished the hatchability of the fertilized eggs without impacting egg production, suggesting that these genes’ activity is essential in egg activation and/or embryogenesis but not for oogenesis (Table 1, Figure 1A, Table S1). For the nine genes in Class 2, depletion of their products led to slight reduction of egg production, but did not significantly influence the hatchability of the eggs. These genes may be involved in oogenesis, but their maternally-derived products are not essential thereafter. It is also possible that their expression was not sufficiently knocked down to fully block oogenesis (or to show later effects).

**Figure 1.**
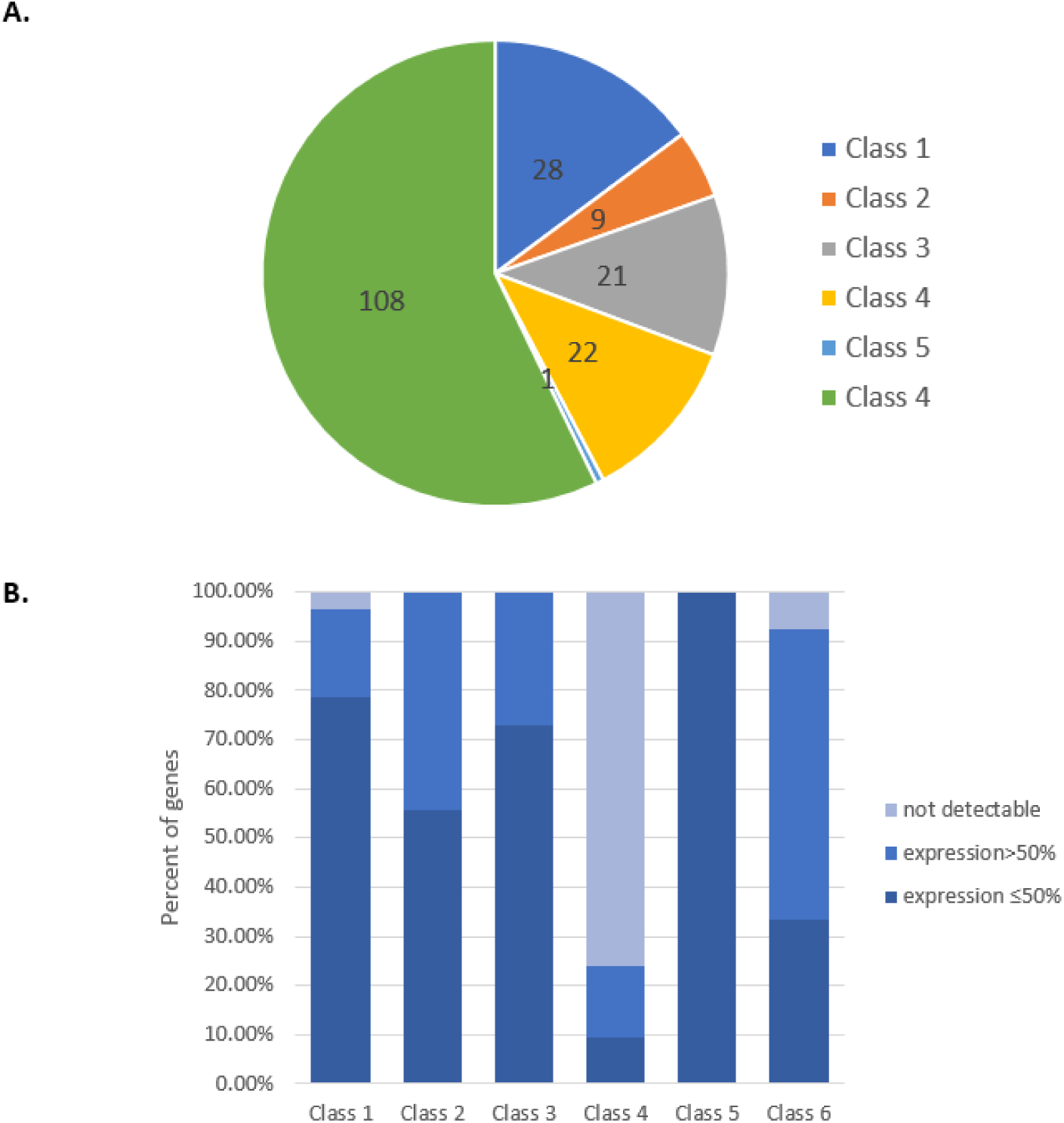
**Germline-specific RNAi knockdown of maternal proteins that are phospho-regulated upon egg activation exhibited different impacts on female fertility.** (A) The frequency of various fertility phenotypes observed in RNAi females (MTD-Gal4/UAS-shRNA or matαrub-Gal4/UAS-shRNA) crossed to ORP2 males. A total of 189 genes were screened. (B) Knockdown efficiency of the target gene in different phenotype classes. The residue expression of the target genes in the oocytes produced by RNAi females was examined using RT-PCR and band intensity quantification. The expression of genes cannot be evaluated when the knockdown led to abolished egg production due to the lack of oocyte. The expression of 9 genes cannot be detected with RT-PCR.

**Table 1.**
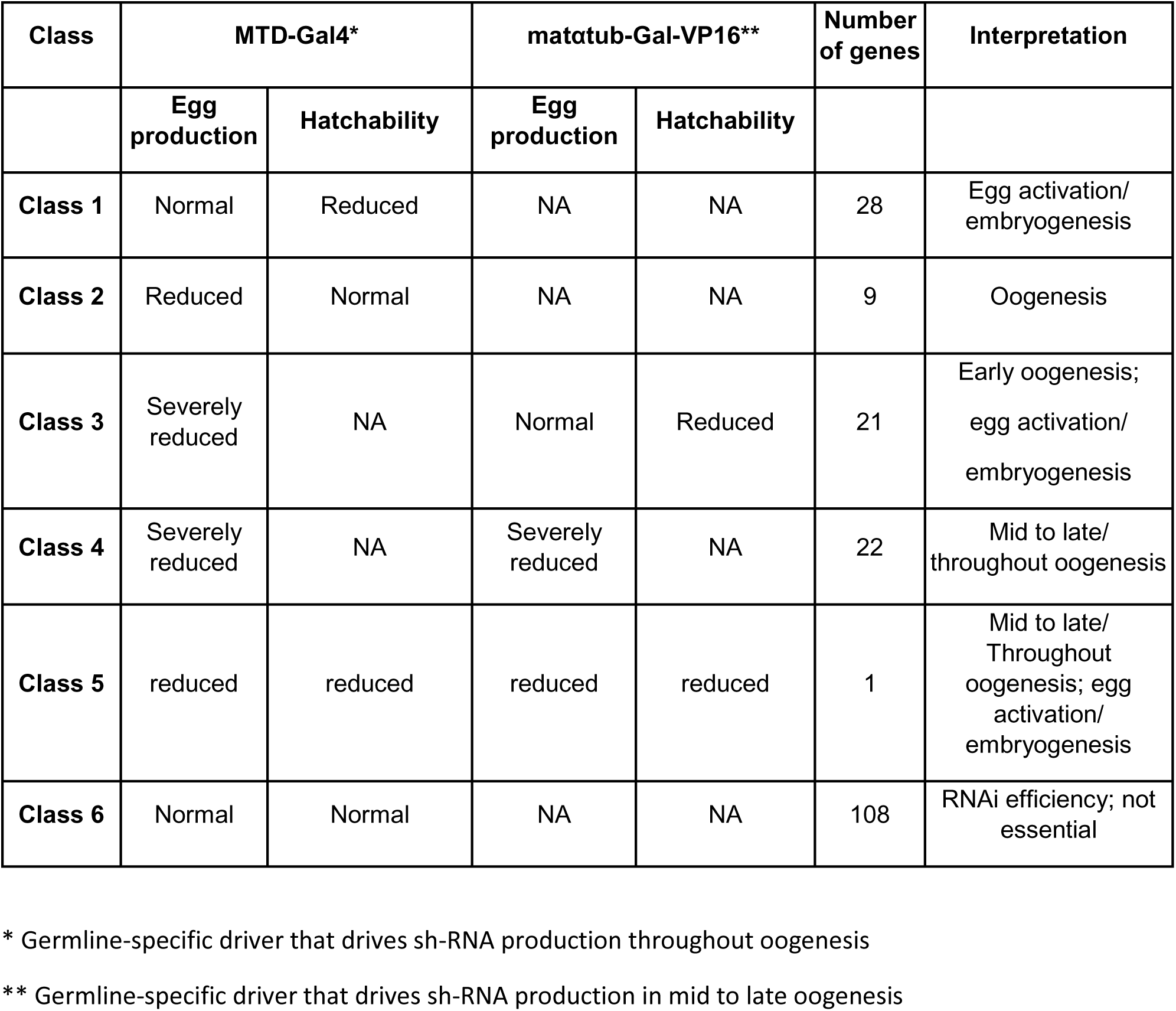
CLASSIFICATION OF FERTILITY PHENOTYPES IN THE PRIMARY AND SECONDARY GERMLINE SPECIFIC RNAi SCREEN.

Genes in Class 3 and Class 4 are likely crucial for oogenesis since the knockdown of their expression throughout oogenesis led to severe reduction or complete abolishment of egg production. For the 21 genes in Class 3, restricting the depletion of their gene products to mid and late oogenesis, by driving knockdown with the matα4-GAL-VP16 driver, circumvented the knockdowns’ impacts on egg production and revealed various hatchability phenotypes. This subset of genes is thus likely important in early oogenesis, as well as in egg activation or embryogenesis, but their expression may not be essential in mid and late oogenesis. However, we cannot rule out the possibility that perdurance of protein products from early stages of oogenesis masked any defects from the knockdown in later stages of oogenesis, or that knockdown of these genes driven by *matα4-GAL-VP16* was insufficient to generate an oogenesis phenotype. For the 22 genes in Class 4, temporal manipulation of their knockdowns did not alter the impact of these knockdowns on egg production. Thus expression of these genes is likely required throughout oogenesis for normal egg production, and it is impossible to determine whether they also affect egg activation or early embryogenesis. Knockdown of Acn significantly impacted both egg production and hatchability when driven by either driver, indicating its involvement in late oogenesis as well as egg activation or embryogenesis. Hence, we assigned Acn to a different class, Class 5.

For the 108 genes in Class 6, the knockdown of their expression throughout oogenesis did not impact female fertility. RT-PCR indicated low RNAi efficiency of < 50% in 59% of these knockdowns (Figure 1B), suggesting that the currently available TRiP lines did not permit full evaluation of their roles in oogenesis or early embryogenesis. Ovary expression of eight genes was too low to be detectable under our RT-PCR conditions, making it difficult to access their knockdown efficiency. For the genes that are knocked down by more than 50%, the lack of phenotype means that they might not be uniquely essential for female fertility, but we cannot rule out that the residual gene product was sufficient for their function.

To assess the effect of off-targets in our results, we obtained all available shRNA lines targeting a given gene. Of the 16 sets of shRNA lines that targeted the same genes, 11 generated the same phenotype in all the lines in the set (Table S1). For the other 5 sets of shRNA lines, fertility phenotypes were observed in some lines but not the others in the set. For four of these five sets, the differences in phenotypes are most likely due to differences in knockdown efficiencies, since the lines with no fertility defects showed highest level of residual expression. The two shRNA lines targeting *nap1* generated similar levels of knockdown, thus the difference in phenotypes may be a result of small differences in knockdown that were undetectable by our semi-quantitative methods, to off target effects of one of the constructs, or to unknown genetic or environmental factors that varied between the lines and their tests.

Taken together, the combination of MTD-Gal4 and *matα4-GAL-VP16* driven germline specific RNAi identified 53 genes with possible roles in oogenesis. This group includes factors that are known to mediate essential processes in oogenesis, such as *14-3-3ε (Benton et al 2002)*, *no child left behind (nclb) (Casper et al 2011)*, CG9556 (Pan et al 2014) and *short stop (shot)* (Roper & Brown 2004), validating the effectiveness of our method at capturing relevant genes. Intriguingly, 32 of the 53 genes had no previously reported roles in oogenesis. These genes likely represent new actors in oogenesis that merit further investigation. 50 genes are likely involved in egg activation and/or embryogenesis. Notably, 22 of these 50 genes have no previously characterized maternal phenotypes.

### Maternal knockdowns that led to severe hatchability defects caused developmental arrest at various stages

To investigate the roles of the maternal proteins whose knockdowns caused hatchability phenotypes (Class 1, Class 3 and Class 5), we examined the phenotypes of 2-4-hr old embryos produced by RNAi females fertilized by wildtype (ORP2) males. Genes whose knockdown reduced the hatchability of the embryos to 50% or below were prioritized (43 genes).

*Drosophila melanogaster* embryos go through 14 syncytial mitotic divisions (embryogenesis stage 1-4) in the first 2 hrs of development, before cellularization takes place at stage 5 of embryogenesis. Embryonic development before cellularization is largely maternally driven (Campos-Ortega & Hartenstein 1997). At 2-4 hrs post fertilization, more than 80% of fertilized embryos produced by control females developed to stage 4 or later stages of embryogenesis (Figure 2A, Table S3), while in the knockdown embryos, we observed developmental arrest in various stages of embryogenesis (Figure S2, Table S3). We were especially interested in knockdowns that led to arrest at stage 1 (apposition of male and female pronuclei) and 2 (syncytial mitotic division 1-8) of embryogenesis, since early arrest likely reflects defects in egg activation or the transition into zygotic syncytial divisions.

**Figure 2.**
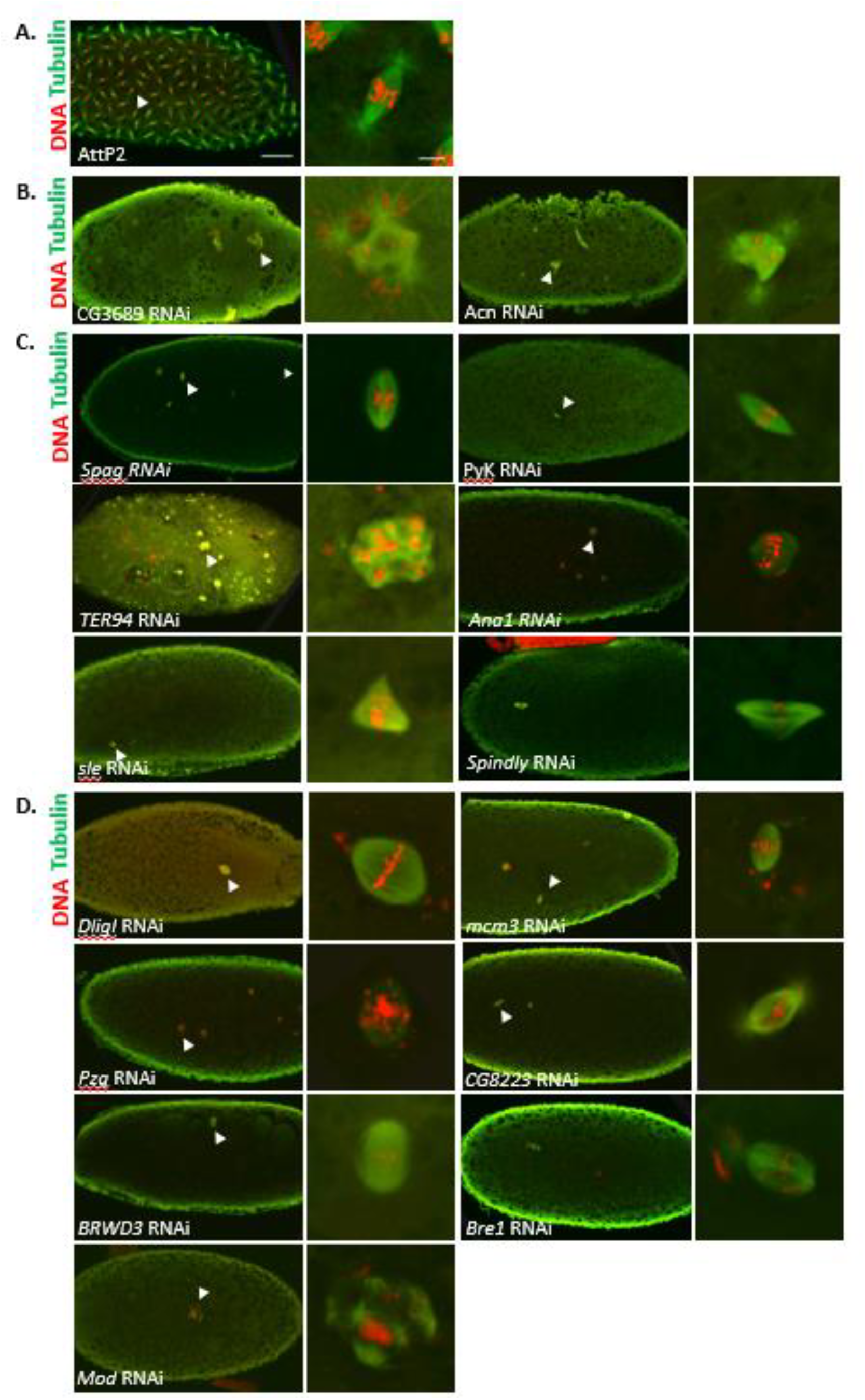
**Maternal knockdown of 15 genes led to developmental arrest in stage 1 and/or early stage 2 of embryogenesis.** 2-4 hr old embryos from knockdown females, stained for tubulin (green) and DNA (red). For each set of images, the one on the right shows a close-up view of the spindle (marked by the arrow on the left). (A) embryos from AttP2 control females mated with ORP2 males. (B) embryos from *CG3689*, *Acn* knockdown females arrested in stage 1 and early stage 2 of development and exhibiting abnormal mitotic spindle organization. (C) embryos from *spag, PyK*, *TER94*, *Ana1*, *sle* and *spindly* knockdown females arrested in stage 1 and early stage 2 of development and exhibiting abnormal centrosome and spindle organization. (D) embryos from *DligI*, *Mcm3*, *Pzg*, *CG8223*, *BRWD3*, *Bre1* and *mod* knockdown females arrested in stage 1 and early stage 2 of development and exhibiting abnormal chromosome formation and segregation. Scale bars: embryo 50 µm, nucleus 5 µm.

We observed that for 27 genes, maternal knockdown caused a significant portion of the embryos to arrest in stage 1 and/or early stage 2 of embryogenesis. Knockdown of 17 of these genes caused a large proportion of embryos to arrest in stage 1 of embryogenesis, prior to completion of first mitosis: *DNA ligase I* (*Dlig-I*) (77%, n = 36)*, minichromosome maintenance* (*mcm3*) (80%, n = 25)*, spindly* (54%, n = 26), *spc105r* (91%, n = 34), *ballchen* (*ball*) (89%, n = 28), *BRWD3* (38%, n = 60), *mbs* (40%, n = 30), *mri* (87%, n = 31), *pyruvate kinase* (*pyk*) (46%, n = 48), *bre1* (43%, n = 35), *mod(mdg4)* (58%, n = 26), *slender lobes* (*sle*) (30%, n = 27), *aubergine* (*aub*) (68%, n = 28), *αTub67C* (96%, n = 25), *acn* (36%, n = 28), *spaghetti (spag)* (13%, n = 72) *and modulo* (*mod*) (41%, n = 17) (Table S3). These phenotypes may indicate roles of these genes in egg activation or in initiation of syncytial division. Maternal knockdown of 17 of these genes caused significant embryogenesis arrest in early stage 2: *TER94* (86%, n = 21), *spindly* (38%, n = 26)*, 14-3-3ε* (86%, n = 21), *pzg* (91%, n = 22), *mbs* (40%, n = 30), *su(var)205* (40%, n = 15), *PyK* (52%, n = 48)*, mod(mdg4)* (42%, n = 26), *aub* (32%, n = 28), *sle* (63%, n = 27), *MCPH1* (87%, n = 23), *ana1* (100%, n = 20), *pk92B* (42%, n = 33), *plutonium* (*plu*) (71%, n = 21), *spag* (19%, n = 72), *CG8223* (61%, n = 33), *CG3689* (40%, n = 48). Note that the knockdown of seven genes: *mbs*, *sle*, *mod(mdg4)*, *spag*, *spindly*, *PyK* and *aub*, led to significant arrest in both stage 1 and stage 2 of embryogenesis; thus they are included in both categories here. In addition, the knockdown of *eco* caused significant developmental arrest in late stage 2 (63%) of embryogenesis

We confirmed that the lack of hatchability was not due to fertilization failure, by testing for the presence of a sperm tail in 0.5-1.5-hr old embryos produced by knockdown females (Karr 1991). We found no outstanding reduction in fertilization rate in any of the knockdown embryos (Figure S3, Table S4). Examination of the sequences of the proteins encoded by these 27 maternal-effect genes revealed that many encode proteins with consensus phosphorylation sites for conserved kinases that are involved in or modulated during *Drosophila* egg activation, including Cdk1 (deactivated upon egg activation, (Swan & Schupbach 2007)), Erk (activity decreases upon egg activation, (Sackton et al 2007)) and GSK3 (activity required for egg activation in Drosophila (Takeo et al 2012)) (Table S5). The presence of these consensus sites suggests that these proteins may be targets of these kinases, consistent with their identification as phospho-modulated during this transition, and suggesting the functional importance of their phospho-regulation at this time. Several of the proteins also carry consensus sites for CaMKIIγ, the calcium dependent kinase that plays essential role in vertebrate egg activation, but has not yet been tested for involvement in Drosophila egg activation.

The 27 genes include 12 with previously characterized maternal-effect phenotypes, validating the premise of our screen that the phospho-regulated proteins include ones whose action is essential during this transition. These genes include *ball* (Ivanovska et al 2005, Nikalayevich & Ohkura 2015), *plu* (Elfring et al 1997, Shamanski & Orr-Weaver 1991)*, mod(mdg4)* (Buchner et al 2000)*, spc105r* (Radford et al 2015, Yan et al 2014)*, mri* (Krauchunas et al 2012)*, su(var)205* (Kellum & Alberts 1995), *pk92B* (Sopko et al 2014), *αtub67C* (Komma & Endow 1997, Matthews et al 1993), *mcph1* (Brunk et al 2007), *mbs* (Sopko et al 2014), *aub* (Mani et al 2014), and 14-3-3ε (Perrimon et al 1996). In most cases, the knockdown phenotypes we observed recapitulated the previously reported maternal-effect phenotypes; the few small discrepancies are likely due to incomplete RNAi efficiency. As two examples, *ball* and *plu* are known to be essential for a proper egg-to-embryo transition. *ball* encodes a kinase that is crucial for the condensation of chromosomes during meiosis (Nikalayevich & Ohkura 2015). We observed that Ball depletion in the female germline caused 89% of her embryos to arrest before completing their first mitosis, with aberrant chromosome arrangements in the polar body. This resembles the previously reported phenotype observed in embryos produced by *ball* mutant females (Ivanovska et al 2005, Nikalayevich & Ohkura 2015). 11% of *ball*-knockdown embryos completed the first mitosis and arrested later, in stage 2; given the reported mutant phenotype, we believe the progression of these few embryos reflects incomplete knockdown in the egg chambers that gave rise to them. *plu* encodes a component of Pan Gu Kinase (PNG), which controls the translation of important maternal transcripts such as *smg* and *cyclin B* upon egg activation (Elfring et al 1997, Shamanski & Orr-Weaver 1991). *plu* mutant females produce embryos that arrest in early embryogenesis with giant polyploidy nuclei as a result of mis-regulation of DNA replication and failure of mitosis. We observed the same phenotype in all *plu* knockdown embryos (Figure S2, Table S3, Table S4). Our recapitulation of phenotypes reported in previous studies using germline clones, RNAi, or hypomorphs validated the effectiveness of our methods in detecting genes with maternal-effect phenotypes. A full summary of the phenotypes observed in maternally depleted embryos for the 27 genes can be found in Table S3.

Intriguingly, for 15 of the 27 genes no maternal-effect phenotype had previously been reported. We discuss these genes in more detail below. The early arrest embryos resulting from maternal knockdown of these 15 genes exhibited 3 distinct categories of abnormalities, including abnormal mitotic spindles, abnormal centrioles and abnormal chromosome morphology.

#### Abnormal mitotic spindles

For the two genes in this category, their maternal knockdown led to mitotic spindles with abnormal morphology and multi-polar spindles in the early embryos, likely due to mis-regulation of centriole amplification or aberrant spindle organization. Maternal knockdown of Acn led to 36% stage 1 arrest and 29% stage 2 arrest among the embryos (Table S3). Metaphase spindles with abnormal shapes or multi-polar spindles were observed in 17% of the early arrested *acn* knockdown embryos (Figure 2B). Acn is important for endosomal trafficking and the stabilization of early endosomes (Haberman et al 2010). It is also involved in RNA splicing as an accessory factor of the exon junction complex (Hayashi et al 2014). These fundamental cellular roles may explain the slight reduction of egg production that we observed in *acn* knockdown females. Our results indicate that *acn* is a maternal-effect gene whose product is required for the initiation of embryogenesis. It is also possible that the abnormal spindle phenotype in *acn* knockdown embryos is a result of incorrect processing of Acn target mRNAs.

Maternal knockdown of *CG3689* led to 21% stage 1 arrest and 40% stage 2 arrest among the embryos (Table S3). Strikingly, misshapen or multi-polar metaphase spindles were observed in 51% of early arrest embryos with *CG3689* depletion (Figure 2B). Intriguingly, CG3689 is the fly ortholog of mammalian NUDT21, which binds to specific sequences at 3’ end of pre-mRNAs and plays an important role in mRNA processing (Dettwiler et al 2004, Ruegsegger et al 1998, Yang et al 2010). Previous work in Drosophila indicates widespread polyadenylation of maternal mRNAs during egg activation. This polyadenylation process is known to be partially mediated by GLD-2 class poly-(A) polymerase Wispy (Benoit et al 2008, Cui et al 2008, Cui et al 2013). It will be interesting to explore whether CG3689 is involved in the polyadenylation of maternal transcripts upon egg activation and whether it works together with Wispy.

#### Abnormal centrioles

In the embryos from knockdown females for the 6 genes in this category, centrioles were absent from the spindle poles and localized abnormally in the cytoplasm. Maternal depletion of *PyK* led to 46% stage 1 arrest and 52% stage 2 arrest among the embryos (Table S3). Interestingly, centrioles were undetectable at the spindle poles of metaphase nuclei in essentially all of the early arrest *PyK* knockdown embryos. We found many embryos with metaphase spindles that resembled meiotic spindles in shape although they were usually not located at the periphery of the cell, where the meiotic spindles normally are located. We did not observe polar bodies in most of these embryos, suggesting that they indeed failed to complete meiosis. *PyK* encodes pyruvate kinase, an important component of the glycolytic pathway. Our results suggested that glycolysis may be actively regulated during egg activation. It is possible that this regulation is functionally important for meiosis completion.

Maternal depletion of *Spaghetti (Spag)* led to 13% stage 1 arrested and 19% stage 2 arrested embryos (Table S3). Similar to the case of *PyK* knockdown, we found around 48% of the early arrested *Spag* knockdown embryos lacked centrioles at the spindle poles, and had metaphase spindles that resemble the shape of meiotic spindles (Figure 2C, Table S3). However, unlike the case in *PyK* knockdown embryos, polar bodies were present in most of the early arrested *Spag* knockdown embryos, indicating meiosis completion. *Spag* is a regulatory co-chaperone of Hsp70 and Hsp90 that is potentially important for stabilization and assembly of RNA polymerase II complex (Benbahouche Nel et al 2014). Our results indicate that the presence of Spag is necessary for the transition from oocyte to early embryo, but its role in this process is not yet known.

Spindly is a highly conserved protein that is required for recruitment of dynein to kinetochores during cell division. It is known to be crucial for the metaphase to anaphase transition in mitosis in Drosophila (Griffis et al 2007) and essential for oocyte maturation in mouse (Zhang et al 2010). Maternal-effects had not previously been reported for *spindly*. We found that knockdown of *spindly* in oocytes led to metaphase arrest before the completion of the second mitotic division (Table S3). The polar bodies in 63% of the knockdown embryos exhibited abnormal microtubule and chromosome arrangements, consistent with *spindly’s* reported function as a dynein recruiting factor (Griffis et al 2007). Our results indicate that maternal deposition of Spindly is crucial for the progression of mitotic divisions in early embryos.

Significant fractions of embryos maternally depleted for *TER94* (10% stage 1, 86% early stage 2), *sle* (30% stage 1, 67% stage 2), and *ana1* (100% early stage 2) also arrested early in embryogenesis (Table S3). Most *TER94 or sle* knockdown embryos and all *ana1* knockdown embryos arrested at stage 2 of embryogenesis. Strikingly, the M phase nuclei in the *TER94, sle* and *ana1* knockdown embryos generally showed abnormal spindle morphologies. Most ana1 knockdown embryos lacked centrioles at the spindle poles, while in most *TER94* and *sle* knockdown embryos, centrioles frequently aggregated or localized abnormally and detached from M phase spindles (Figure 2C, Table S3). In addition, chromosome number and morphology were abnormal in M phase nuclei in some *TER94* and *sle* knockdown embryos. As TER94 and Ana1 are known to be crucial for microtubule organization and centrosome formation respectively, their roles in the facilitation of early syncytial divisions are interesting but not surprising. On the other hand, Sle is known to be involved in nucleolar organization and rRNA processing (Orihara-Ono et al 2005). Our results revealed a novel function for Sle, as a maternal protein crucial for early embryogenesis.

#### Abnormal chromosome morphology and segregation

For the seven genes in this category, their maternal knockdown led to abnormal morphology and segregation of the chromosomes in early embryos. Maternal knockdown of *mcm3* (80% stage 1, 20% stage 2, n = 25), *Dlig-I* (83% stage 1, 17% stage 2, n = 36)*, CG8223* (9% stage 1, 61% stage 2, n = 33)*, pzg* (5% stage 1, 91% stage 2, n = 22)*, Bre1* (43% stage 1, 29% stage 2, n = 35), *mod* (41% stage 1, 29% stage 2, n = 17) and *BRWD3* (38% stage 1, 13% stage 2, n = 60) caused large proportions of embryos to arrest early in embryogenesis (Table S3) with prevalent abnormalities in the morphology and segregation of chromosomes, and a range of other defects in the embryos produced by females depleted of the products of these genes (Figure 2D). In 28% of *Dlig-I* knockdown embryos and all *pzg* knockdown embryos, clusters of small, short chromosomes were observed, indicating possible chromosome disintegration. In *mcm3, bre1,* and *CG8223* knockdown embryos, chromatin strings were frequently observed extending from the chromosomes as they lined up on the metaphase plate (Figure 2D), which may indicate aberrant chromosome organization or chromosome disintegration. In *mod* knockdown embryos, we frequently observed aggregation of chromatin that seemed unable to segregate properly, suggesting a failure of chromosome formation and segregation. 20% of early arrested BRWD3 knockdown embryos had metaphase nuclei with small condensed chromosomes (Figure 2D), indicating a possible defect in chromosome condensation dynamics.

*mcm3, Dlig-I and pzg* are essential genes for DNA replication and/or DNA damage repair, which explains their importance for the progress of syncytial divisions and chromosome integrity, but this is the first report of their maternal-effects in Drosophila. Mod is crucial for the assembly of centromeres in mitosis and is also essential for meiosis and spermatid differentiation in males (Mikhaylova et al 2006). Maternal depletion of Mod does not seem to affect meiosis in females since we observed the presence of polar bodies in 95% of *Mod* maternal knockdown embryos (Table S3). However, our results indicate that maternally-deposited Mod is essential for progression of embryonic mitosis. Bre1 was shown to be important for the maintenance of germline stem cells (Xuan et al 2013). However, in addition to confirming the oogenesis defects reported for Bre1 mutants in previous studies, our results uncovered a need for Bre1 in chromosome organization in meiosis or early syncytial divisions. BRWD3 is known to negatively regulate histone H3.3 deposition to the chromatin by interacting with HIRA complex and its associated chaperon Yemanuclein (Yem) in Drosophila (Chen et al 2015). Maternal histone H3.3 is assembled on paternally derived chromosomes shortly after fertilization (Horard & Loppin 2015, Loppin et al 2005). This process is crucial for the subsequent combination of male and female genetic material and is known to be mediated by HIRA (Loppin et al 2005). It is possible that BRWD3 is involved in the fine-tuning of the maternal H3.3 deposition, so the histone variant is assembled onto the male-derived chromosomes.

Seventy percent of embryos with maternal depletion of a novel protein CG8223 arrested by early stage 2 of embryogenesis with abnormal chromosome organization. In metaphase spindles of the knockdown embryos, aberrant strands of chromatin that stray from the chromosomes were frequently observed, indicating that CG8223 may be an as-yet uncharacterized player in chromosome organization and segregation. Consistent with this idea, the mammalian ortholog of CG8223 is involved in the histone deposition that accompanies DNA replication (Alekseev et al 2003, Finn et al 2008, Richardson et al 2006).

In addition to genes whose maternal knockdown caused early embryonic arrest, we also identified 16 genes: *shot*, *RPA2*, *inaD*, *CG9556*, *nipped*-*b*, *AP-2α*, *myopic*, *eco*, *vps2*, *axin*, *msn*, *CG32088*, *rec*, *nucb1*, *D1* and *akap200*, whose maternal knockdown led to developmental arrest at later stages of syncytial divisions and embryogenesis (Table 2, Table S3). Although it is clear that these genes function in later stages of embryogenesis, the possibility of incomplete knockdown masking an early embryonic requirement cannot be ruled out.

**Table 2.** CANDIDATES ARE REQUIRED AT DIFFERENT STAGES OF OOGENESIS AND EMBRYOGENESIS.

| Stage 1-2 oogenesis | post stage 2 oogenesis | stage 1-2 embryogenesis | post stage 2 embryogenesis |
| --- | --- | --- | --- |
| TER94 | Rpn9 | TER94 | shot |
| shot | scny | CG5602 | RPA2 |
| Dlig-I | Hel25E | mcm3 | inaD |
| mcm3 | Prosa7 | Su(var)205 | CG9556 |
| 14-3-3ε | nclb | pzg | Nipped-B |
| ballchen | CG2263 | spag | AP2α |
| spindly | cct5 | BRWD3 | myopic |
| spc105r | lolal | spindly | eco |
| CG2915 | Tcp1eta | spc105r | Vps2 |
| Su(var)205 | rtf1 | ballchen | axin |
| pzg | Uba2 | mrityu | msn |
| spag | Tango7 | PyK | CG32088 |
| RPA2 | CG31368 | Bre1 | rec |
| CG9609 | NAT1 | mod(mdg4) | NUCB1 |
| BRWD3 | eIF5B | aub | D1 |
| Upl1 | eIF2α | slender lobe | Akap200 |
| inaD | Uba1 | MCPH1 |  |
| CG9556 | Rpl22 | αTub67C |  |
| D1 | eIF4a | ana1 |  |
| bre1 | CDK12 | Pk92B |  |
| Mbs | Acn | modulo |  |
|  | eIF2Bσ | Plutonium |  |
|  |  | CG8223 |  |
|  |  | CG3689 |  |
|  |  | Acn |  |
|  |  | 14-3-3e |  |
|  |  | Mbs |  |

### Several maternal protein knockdowns that lead to early developmental arrest also induced defects in egg activation events

Early embryogenesis arrest may reflect defects in egg activation or the transition into zygotic syncytial divisions. To investigate whether the maternal knockdowns that caused early developmental arrest (27 genes) had any impacts on egg activation, we examined 3 major events of egg activation in the embryos produced by the knockdown females, including changes in egg covering, translation initiation and meiosis completion.

The resumption and completion of meiosis is an important aspect of egg activation. In *Drosophila melanogaster*, one of the meiotic products becomes the female pronucleus upon the completion of meiosis while the other meiotic products combine and form a polar body with distinct morphological traits (condensed chromosomes surrounded by microtubules) (Mahowald et al 1983). We interpreted the presence of a normal polar body as a signature of meiosis completion, in 0.5-1.5 hour old embryos (Table S4). Normal polar bodies were observed in 93% of the embryos produced by control (AttP2) females (Figure 3). We found no major aberrancy in polar body presence and morphology in maternal knockdown embryos of 15 genes (Table S4). However, we found an absence of polar bodies in large fractions of embryos from females with germline knockdown of *PyK* (65% absent, n = 31), *mri* (38% absent, n = 37), *Acn* (33% absent, n = 18), *BRWD3* (30% absent, n = 23) and *pk92B* (36% absent, n = 28), suggesting defects in meiosis completion in these embryos. The absence of polar bodies in a large fraction of PyK knockdown embryos suggests that the abnormal spindle we observed in 2-4 hr old PyK knockdown embryos may indeed have been meiotic spindles, thus further confirming that meiosis is not completed in these embryos. We also found prominent abnormalities in the morphology of polar bodies in embryos with maternal knockdown of several genes (Figure 3), including *Dlig-I* (83% abnormal, n = 23), *mcm3* (56% abnormal, n = 36), spc105r (96%, n = 23), plu (100%, n=14), *spindly* (63% abnormal, n = 19), aub (57%, n = 28), *mri* (62% abnormal, n = 37), *14-3-3ε* (93% abnormal, n = 29), *mod(mdg4)* (58% abnormal, n = 24), ball (88%, n = 24) and PyK (13%, n = 31), indicating defects in meiosis or chromosome condensation and microtubules organization that are involved during the formation of polar bodies in these knockdown embryos.

**Figure 3.**
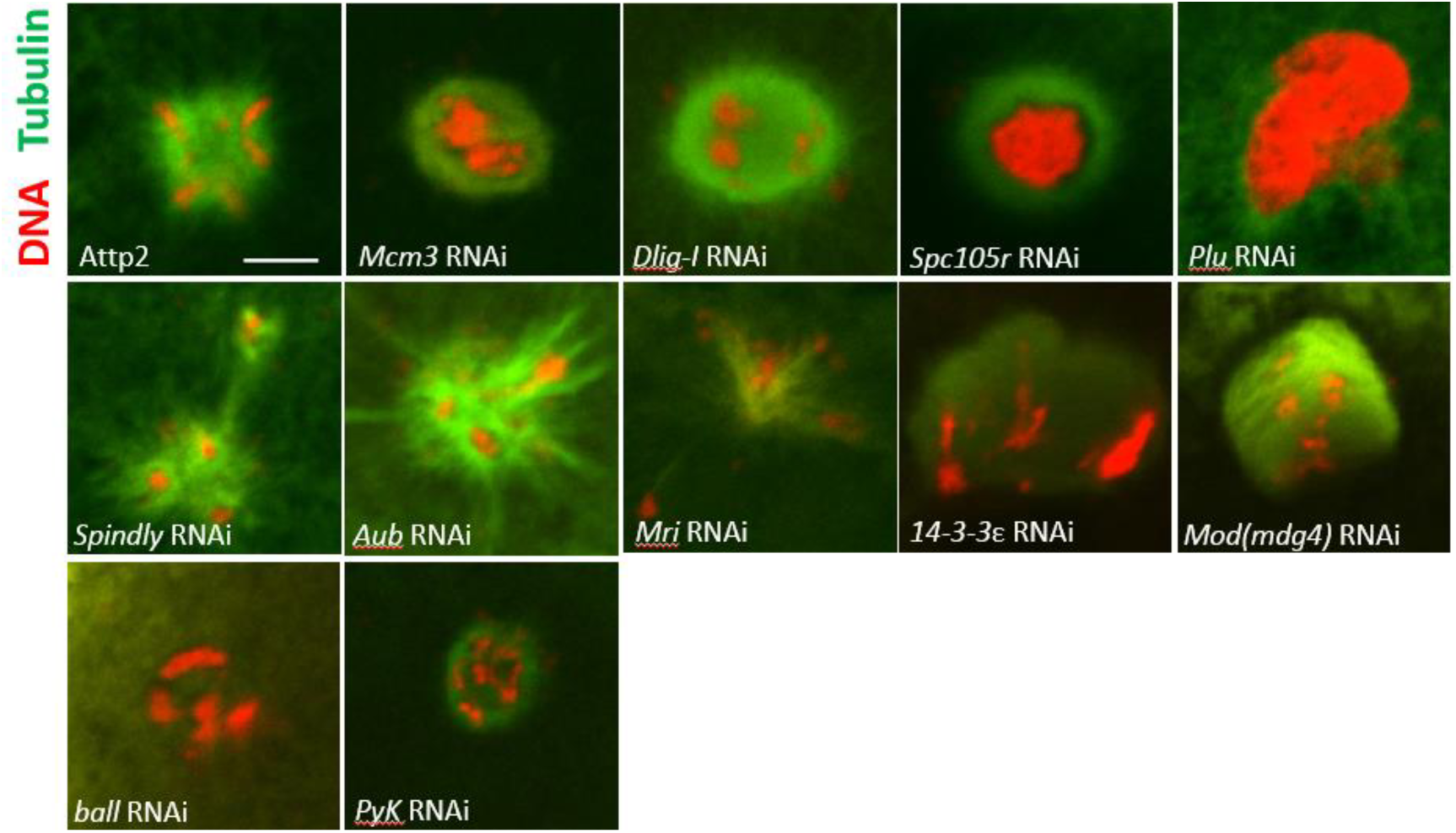
**Polar bodies with abnormal morphology were observed in knockdown embryos of 11 genes.** Significant portions of 0.5-1 hr embryos with maternal knockdown of Dlig-I (83% abnormal, n=23), mcm3 (56% abnormal, n=36), spc105r (96%, n=23), plu (100%, n=14), spindly (63% abnormal, n=19), aub (57%, n=28), mri (62% abnormal, n=37), 14-3-3ε (93% abnormal, n=29), mod(mdg4) (58% abnormal, n=24), ball (88%, n=24) and PyK (13%, n=31) had polar bodies with abnormal morphology. Tubulin is in green. DNA is in red. Scale bars: 5µm

In *Drosophila melanogaster*, egg activation includes the cross-linking of the vitelline membrane outer layer and the chorion, causing the egg coverings to become impermeable to small molecules (Heifetz et al 2001). We treated the 0.5-1.5 hour old eggs produced by knockdown females with 50% bleach to examine this biochemical change of the egg coverings. We found that the maternal knockdown of *acn* caused a large reduction in bleach resistance; only 41% of the eggs produced by the knockdown females persisted longer than 2 min in 50% bleach (Table S4), suggesting a role of Acn in facilitating eggshell production or triggering the cross-linking of egg covering. Maternal knockdown of the other 26 genes did not significantly affect the bleach resistance of the eggs.

Egg activation is also accompanied by a dramatic increase in the translation of many maternal mRNAs important for subsequent embryogenesis (Horner 2007, Kronja et al 2014), including (in Drosophila) *smg, bcd, zld* (Driever & Nusslein-Volhard 1988, Eichhorn et al 2016, Tadros et al 2007). We used western blots to examine the translation of *smg* in 0-1hr activated but unfertilized eggs produced by knockdown females, as a marker of a general increase in translation. As expected, Smg protein is detected in the lysate of activated eggs but not in that of stage 14 oocytes produced by control females (Tadros et al 2007). Consistent with previous findings, Smg protein was not detected in activated eggs produced by females with germline depletion of Plu, which is a subunit of Pan Gu Kinase, and essential for the translation of Smg upon egg activation (Tadros et al 2007). We found no significant disturbance in *smg* translation in eggs with any other maternal knockdowns, indicating that none of the genes examined, except for Plu, are essential for the general translational increase that occurs upon egg activation or the translation of *smg* specifically (Figure S4). We cannot rule out the possibility that the 26 genes whose knockdown did not affect *smg* translation may include factors that regulate the translation of specific maternal mRNAs other than *smg*.

## DISCUSSION

Although screens for maternal-effect mutations have identified some genes needed during the egg-to-embryo transition, the number of genes that they have uncovered has been surprisingly small. This suggests that other genes whose function is critical at this time may be essential later in development, making it impossible to detect them in whole-organism screens for maternal-effect mutations. Alternative methods are needed to find such genes. Although germline-specific depletion can identify phenotypes for such genes, efficient screening requires enrichment for candidates. One type of enrichment has been to focus on genes with enriched germline expression. An RNAi screen among genes of this type in *C. elegans* revealed that 322 out of 766 ovarian enriched genes are required for normal egg production and/or embryogenesis (Piano et al 2002). A similar screen in Drosophila found that 10.5% of the 3491 germline-enriched genes were important for female fertility (Yan et al 2014). However, since not all genes that are important for female fertility are preferentially expressed in the female germline, functional screen of germline-enriched genes may fail to capture important regulators of female fertility that do not have enriched germline expression. Our study addressed this potential problem by searching among maternal proteins that are phospho-regulated during egg activation. Out of the 189 candidate genes, we identified 81 that are crucial for different aspects of female fertility, indicating that this set of phospho-regulated maternal proteins is enriched for important modulators of oogenesis and early development.

Intriguingly, 88% of the genes we identified as needed for these processes have homologs in mouse. The mammalian orthologs of seven of the genes: *ball* (mouse *vrk1*) (Wiebe et al 2010), *MCPH1* (mouse *mcph1*) (Liang et al 2010), *14-3-3ε* (mouse *ywhae*) (Kosaka et al 2012), *rec* (mouse *mcm8*) (Lutzmann et al 2012), RhoGap68F (mouse Arhgap1) (Wang et al 2005), *spindly* (mouse *spdl1*) (Zhang et al 2010) and *CG9556* (mouse *cops2*) (Lykke-Andersen et al 2003) have previously been reported to be important for female fertility in mouse, suggesting conserved roles of these genes in female reproduction. For the vast majority of the fly fertility factors uncovered by our screens the roles of their orthologs in female fertility have not yet been examined in other organisms. The conservation of these genes, and make them good candidates for investigation as regulators of egg activation or early embryogenesis in organisms beyond Drosophila.

The temporal control of germline specific knockdown and examination of arrest stages in the knockdown embryos identified approximate stages of oogenesis and embryogenesis that required the activity of each gene. The functional classes enriched in these subsets of genes corresponded closely with the major developmental events that occur in these stages (Figure S1B). 21 genes are likely crucial for the early events in oogenesis since driving shRNA expression early but not late in oogenesis led to defects in egg production. This group of genes is enriched in factors involved in mitotic cell cycles and regulation of cellular processes (Figure S1B). These categories are consistent with the early events in Drosophila oogenesis that take place in the germarium, which include germ cell differentiation followed by four rounds of mitosis that generate a cyst of 16 germ cells among which one will be specified as oocyte (King et al 1968). Phospho-regulation of these proteins during egg activation is particularly interesting since they are likely not critical for later stages of oogenesis, but play essential roles in early embryogenesis. It is possible that these candidates are inactivated during mid and late oogenesis, and are activated again through phospho-regulation to facilitate the rapid cycles of syncytial division that follows egg activation. 22 genes are likely essential in mid to late or throughout oogenesis since temporal manipulation of shRNA expression did not alter the detrimental impact of the knockdown on egg production. This group of genes is highly enriched in factors regulating translation and the synthesis of other biomolecules, consistent with the synthesis of maternal protein and mRNA thats take place during mid and late oogenesis. These proteins may be phospho-regulated during egg activation to contribute to dynamic regulation of translation at the initiation of embryogenesis, including the termination of oogenesis progression.

Of the 43 genes that caused more than 50% reduction of egg hatchability when knocked down in female germline, 27 are likely important for egg activation and initiation of syncytial divisions since maternal knockdown of these genes led to significant early developmental arrest in the embryos. This set of genes include several that are needed in essential cellular processes including DNA damage repair and replication. Given these functions, it is not surprising that perturbation of their expression in the mature oocyte causes severe defects in embryo development. Their detection among proteins whose phospho-state changes during egg activation validates the hypothesis that led to our screen, and suggests that their activity may be post-translationally controlled and modulated during this transition, something that will bear future molecular study. 16 of the 43 genes are needed for the progression of syncytial divisions and possibly later events in embryogenesis since maternal knockdown led to embryonic arrest at stage 3 or later stages of embryogenesis. Both subsets are enriched with factors that regulate cell cycle progression, which is consistent with the rapid rounds of syncytial divisions that take place in early Drosophila development.

In addition to discovering new roles for known proteins, we found two novel factors, CG3689 and CG8223, that are important for early syncytial divisions. Both proteins are conserved in mammalian systems. The mammalian ortholog or CG3689 is Nudt21, which is known to play a role in the 3’ polyadenylation of pre-mRNAs (Dettwiler et al 2004, Ruegsegger et al 1998, Yang et al 2010). Since CG3689 shares 77% amino acid sequence homology with Nudt21, it may play a similar role in the polyadenylation of maternal transcripts upon egg activation in Drosophila. We frequently observed misshapen multi-polar spindles in *CG3689* knockdown embryos. Interestingly, embryos from females lacking *Wispy*, a Drosophila poly-(A) polymerase that is known to mediate polyadenylation of maternal transcripts upon egg activation, also arrest in early embryogenesis with abnormal spindle formation (Brent et al 2000, Cui et al 2008). Perhaps maternal transcripts of some spindle-associated proteins are regulated by CG3689, and the phenotype that we observe is caused by the perturbation of these spindle-associated proteins. Though our findings indicate that CG3689 maternal knockdown did not significantly impact *smg* translation upon egg activation, it is possible that CG3689 regulates the polyadenylation of a subset of transcripts other than *smg*. This possibility is supported by the finding that Nudt21 binds to RNA at specific sequences (Yang et al 2010). Examination of the global changes in maternal mRNA poly-A tail length upon egg activation in CG3689 depleted background will be informative to elucidate the functions of CG3689.

The mammalian ortholog of CG8223 is NASP (29% amino acid sequence identity), which is, surprisingly, a sperm-specific protein. NASP is a N1/N2 family histone chaperone involved in nucleosome assembly after DNA replication is completed (Finn et al 2008, Osakabe et al 2010, Richardson et al 2006), which hints at interesting possibilities about the roles of *CG8223* in Drosophila. The phenotype we observed in *CG8223* knockdown embryos is consistent with a role in chromosome organization or chromatin assembly. Intriguingly, our finding suggests that this novel protein may be particularly important for early embryogenesis, since the depletion of CG8223 in early oogenesis did not seem to disrupt the formation of 16-cell germline cysts.

For 108 of the genes that we tested, depletion of their products in the female germline did not impact female fertility. In 59% of the cases, the knockdown efficiency was low. Even in the cases of high knockdown efficiency, there could be functional redundancy, preventing accurate assessment of the phenotype. Although in the future one could imagine screens using targeted editing technique like CRISPR/Cas9, at present these techniques are not sufficiently optimized for germline specific gene perturbation.

Early oogenesis and egg activation can be viewed as reciprocal. The former involves the differentiation of germline stem cells into gametes. The latter, on the other hand, involves the transition of a highly specialized reproductive cell, the oocyte, into a totipotent zygote. Intriguingly, we find 13 genes that are involved in early oogenesis and are also important for egg activation or early embryogenesis. Several of these genes encode factors known to be involved in chromatin organization, such as *su(var)205*, *pzg*, *BRWD3*, *ball* and *bre1*. The activity of these factors may need to be regulated appropriately to catalyze the dynamic reorganization of chromatin landscape that accompanies the differentiation and reverse-differentiation at early oogenesis and egg activation respectively. Understanding how phospho-regulation modulates the activities of these factors and what pathways are involved in the phospho-regulation of these factors in response to the calcium signal that initiates egg activation will give valuable information about the mechanism that facilitate the transition between differentiated and totipotent cellular states.

## ACKNOWLEDGEMENTS

We thank National Institute of Health grant (R21-HD072714) for funding this study. We thank Dr. Tim Karr (Kyoto University) and Dr. Howard Lipshitz (University of Toronto) for their kind gifts of anti-sperm tail and anti-smg antibodies. We thank the TRiP consortium at Harvard Medical School (NIH/NIGMS R01-GM084947) for transgenic RNAi fly stocks used in this study. We thank Hawra Al Lawati and Shakela Mitchell for technical assistance during the screening. We thank Drs. Erika Louise Mudrak and Lynn Marie Johnson from Cornell Statistical Consulting Unit (CSCU) for assistance with data analysis, and Drs. Kenneth Kemphues and Michael Goldberg for helpful discussions and comments on the manuscript.

